# Motor learning refines premotor cortical constraint during skilled forelimb behavior

**DOI:** 10.64898/2026.09.24.754238

**Authors:** Di Chen, Fang Jia, Qiyun Huang, Jin Gong, Shuangqi Gao, Chengjie Wan, Ying Guo, Yuanqing Li

## Abstract

Motor learning transforms variable actions into reliable skilled behavior, but how premotor cortex supports this transformation remains unclear. We longitudinally recorded bilateral local field potentials and single-unit activity from the rat rostral forelimb area while animals acquired a single-pellet retrieval task. Here, we show that improved skill was accompanied by reduced, rather than increased, task-related premotor recruitment. Despite this attenuation, firing remained dominated by suppression around pellet retrieval, and stronger pre-retrieval suppression was associated with greater odds of successful performance. Population neural state analysis further showed that successful actions were most strongly coupled to sustained pre-retrieval engagement early in learning, whereas later success occurred with less population-level engagement. These findings reveal premotor cortical constraint as a population-level process that is progressively refined as motor learning transforms variable actions into reliable skilled behavior.

## 1 Introduction

Motor learning enables animals to adapt their actions to environmental demands and acquire skilled behaviors such as reaching for and retrieving food [1, 2]. With practice, initially variable and unreliable actions become increasingly precise and reproducible, improving the reliability with which behavioral goals are achieved [3]. This transformation requires not only the generation of movement, but also the progressive organization and constraint of motor output toward effective solutions [4]. Extensive work has shown that skill acquisition reshapes motor cortical synapses, local circuits, neuronal ensembles and population dynamics [5–8]. Together, these studies provide a detailed account of how cortical circuits closely coupled to movement execution are remodeled during learning. Yet how higher-order cortical control is reorganized to constrain action as skill develops remains less clear.

Higher-order motor cortical areas are well positioned to address this question, contributing to the organization and refined selection of motor output while interacting with primary motor circuits directly involved in execution [9]. In rodents, the motor cortex contains two functionally distinct forelimb representations: the caudal forelimb area (CFA) and the rostral forelimb area (RFA), a premotor-like representation essential for skilled grasping [10–13]. For example, we have shown that targeted modulation of the RFA regulates skilled grasping execution while engaging bilateral motor cortical networks in rats[14]. Recent simultaneous recording and perturbation experiments further support a partial functional hierarchy between RFA and CFA, revealing asymmetric inter-regional influence despite substantial similarity in their movement-related firing patterns [15]. These findings increasingly clarify the functional position of RFA within the cortical motor hierarchy, but not how population-level control within this premotor node is organized as a skilled action is acquired.

Motor-skill learning reorganizes cortical population dynamics and inter-areal interactions, and behaviorally relevant cross-area signals can become increasingly organized with practice [16]. Early acquisition, however, imposes distinct demands on action selection and control because a reliable motor solution has not yet been established [17]. Consistent with this distinction, cortical contributions to motor behavior can differ markedly between skill acquisition and established performance [18]. Premotor control may therefore be organized differently as skilled behavior emerges. Whether premotor population activity becomes progressively integrated into the neural organization of skilled execution or instead becomes more selective as control demands change remains unresolved. Learning-stage differences alone are insufficient to identify control-related neural dynamics; if premotor population activity contributes to skilled control, its expression should also bear a systematic relationship to the outcome of individual actions.

Here, we longitudinally recorded bilateral LFPs and single-unit activity from the RFA while rats acquired a skilled single-pellet retrieval (SPR) task, allowing cortical dynamics to be examined across learning stages, neural scales and behavioral outcomes. Skilled performance improved while broad retrieval-related RFA activity was attenuated, arguing against a simple progressive amplification of motor cortical recruitment. Yet reduced population recruitment did not reflect functional disengagement: stronger pre-retrieval firing suppression was associated with greater odds of successful retrieval. Across learning, the neuronal population recruited into this suppressive response became smaller and reorganized, while sustained pre-retrieval population neural state engagement showed its strongest relationship with successful performance early in learning. Together, these findings support a transition from broad population control during early skill acquisition toward a more restricted and reorganized form of cortical constraint as skilled performance emerges. Motor learning therefore refines, rather than simply strengthens or withdraws, premotor cortical constraint.

## 2 Results

### 2.1 SPR performance improves through progressive spatial refinement of reaching kinematics

To track cortical dynamics during motor skill acquisition, we longitudinally recorded local field potentials (LFPs) and single-unit spiking activity bilaterally from the rostral forelimb area (RFA) while rats performed the skilled single-pellet retrieval (SPR) task across three distinct learning phases: early (P1, week 1), intermediate (P2, week 3), and late (P3, week 5) (Fig. 1a,b). Rats performed the task with the right forelimb; hence, the left and right RFA corresponded to the contralateral and ipsilateral hemispheres, respectively (Fig. 1b). Task trials were aligned to pellet retrieval onset (defined as time 0), segregating reach execution from subsequent consumption or trial failure (Fig. 1c).

**Fig. 1.**
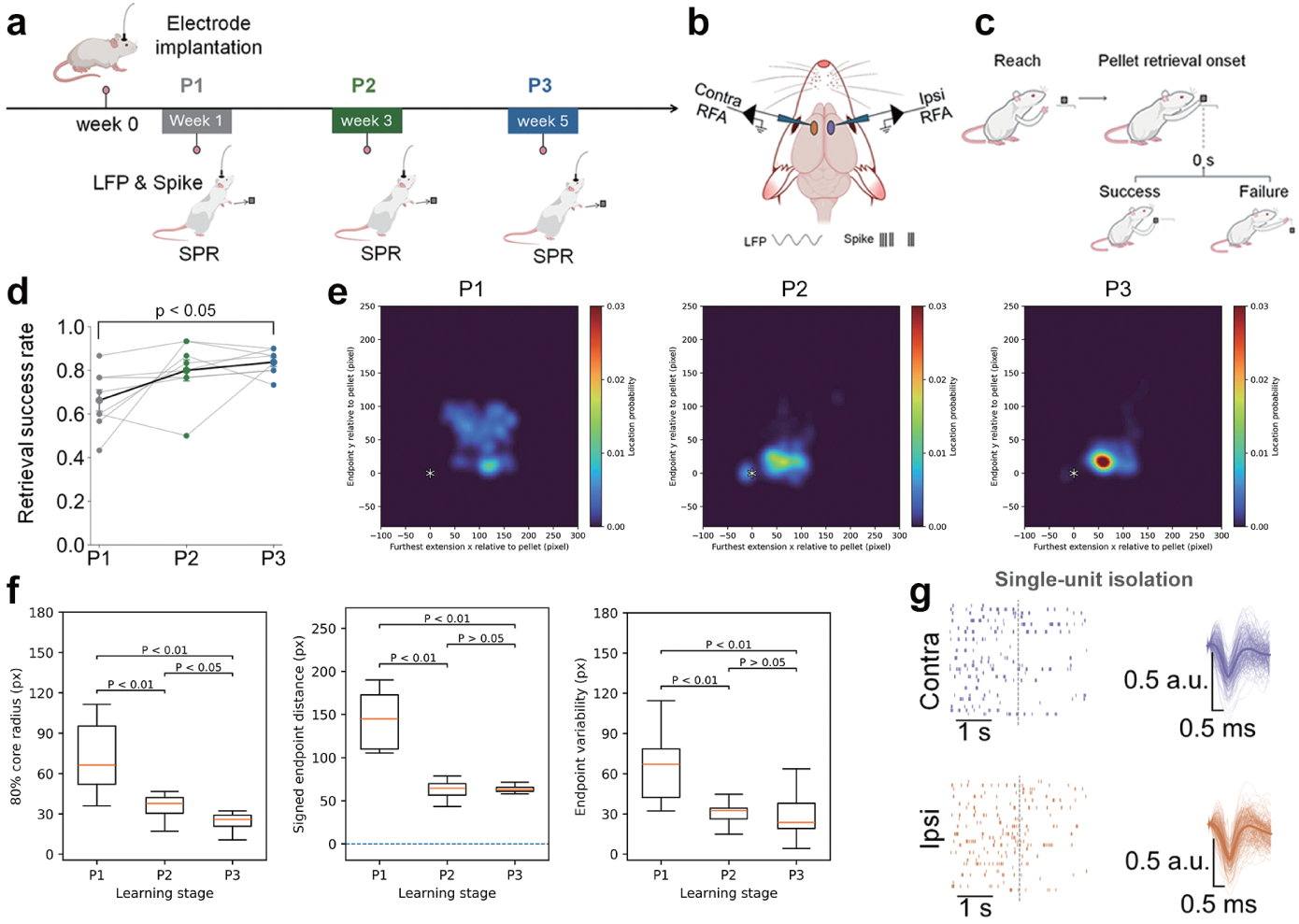
Experimental design and longitudinal bilateral RFA recordings during skilled pellet retrieval learning. a, Experimental timeline. Rats were implanted with recording electrodes at week 0 and performed the SPR task during three learning phases: P1 (week 1), P2 (week 3) and P3 (week 5). Local field potentials (LFPs) and single-unit spiking activity were recorded during SPR performance at each phase. b, Schematic of the bilateral recording configuration in the rostral forelimb area (RFA), showing recording sites contralateral and ipsilateral to the reaching forelimb. c, Schematic of the behavioral sequence during SPR. Pellet retrieval onset was defined as time 0 for subsequent neural analyses. d, Retrieval success rate across learning phases (mean *±* s.e.m., *n* = 8). Lines connect measurements from the same animal and the black line indicates the group mean. P1 versus P2: paired two-sided *t*-test, *t*(7) = *−*2.01, *P*_FDR_ = 0.0849; P1 versus P3: *t*(7) = *−*3.69, *P*_FDR_ = 0.0156. e, Spatial probability density maps of reaching endpoints across learning phases. Endpoints represent paw-tip positions at maximum outward extension relative to the pellet center (asterisk). f, Quantification of endpoint kinematics across stages. Left, 80% core radius (radial boundary encompassing the central 80% of endpoints around the animal centroid). Middle, signed endpoint distance from the pellet along the reaching axis (negative, undershoot; positive, overshoot). Right, endpoint variability. g, Representative single-unit recordings from contralateral (top) and ipsilateral (bottom) RFA, showing trial-aligned spike rasters and normalized spike waveforms. Waveform amplitudes are shown in arbitrary units (a.u.). Scale bars, 1 s for spike rasters and 0.5 ms and 0.5 a.u. for spike waveforms.

Task proficiency improved progressively with training, as retrieval success rate increased from 66.3 ± 4.9% at P1 to 80.0 ± 4.9% at P2 and 83.8 ± 2.0% at P3 (n = 8 rats; Fig. 1d). Performance was significantly improved at P3 relative to P1 (paired two-sided *t*-test, *t*(7) = −3.69, *P*_FDR_ = 0.0156), whereas the increase from P1 to P2 did not reach significance after false discovery rate (FDR) correction (*t*(7) = −2.01, *P*_FDR_ = 0.0849).

We next asked whether this improved proficiency reflected the consolidation of gross task engagement or fine-grained kinematic refinement. Markerless tracking of forepaw trajectories revealed that reaching endpoints became increasingly concentrated around the target across learning (Fig. 1e). Spatial probability density maps demonstrated that the diffuse endpoint distribution characteristic of early acquisition (P1) tightened into a focal cluster adjacent to the pellet at P2 and P3 (Fig. 1e).

Quantitative kinematic analysis corroborated this progressive spatial refinement (Fig. 1f). The 80% core radius—quantifying the spatial dispersion of end-points—shrank markedly from P1 (66.3 ± 5.2 px) to P2 (37.7 ± 3.4 px) and P3 (25.8 ± 2.1 px; Friedman test, P *<* 0.001), with significant reductions across all consecutive stages (Holm-adjusted P *<* 0.05 for P2 vs. P3; P *<* 0.01 for P1 vs. P2 and P1 vs. P3; Fig. 1f, left). Similarly, signed endpoint distance along the reaching axis decreased from 145.0 ± 11.2 px at P1 to 64.4 ± 6.8 px at P2 and 62.8 ± 5.9 px at P3 (P *<* 0.001), reflecting a reduction in initial overshooting that stabilized by P2 (P1 vs. P2 and P3, both P *<* 0.01; P2 vs. P3, P *>* 0.05; Fig. 1f, middle). Reaching endpoint variability likewise dropped significantly by more than half following P1 (P1 vs. P2 and P3, both P *<* 0.01; Fig. 1f, right). Together, these behavioral data demonstrate that skilled motor learning in the SPR task entails a rapid early reduction in spatial variability (P1 to P2) followed by fine-scale asymptotic tuning of movement accuracy (P2 to P3). Chronic recordings across these behavioral stages yielded well-isolated single units with consistent waveform morphologies and trial-aligned modulation across bilateral RFA (Fig. 1g). Thus, this longitudinal paradigm captured the emergence of stereotypic skilled execution, establishing a robust behavioral and electrophysiological foundation to investigate how premotor population dynamics constrain motor output during learning.

### 2.2 Improved SPR is accompanied by attenuated retrieval-related RFA activity

Having established that SPR became more reliable across learning, we next asked how retrieval-related RFA activity was organized as skill developed. Retrieval-aligned event-related potentials (ERPs) showed prominent modulation around pellet retrieval onset during P1 that was attenuated at P2 and P3 in both contralateral and ipsilateral RFA (Fig. 2a,b). Consistent with this pattern, ERP root-mean-square (RMS) amplitude across the full analysis window was reduced at both later learning phases. In contralateral RFA, RMS decreased from 282.7 *±* 50.7 at P1 to 84.2 *±* 13.8 at P2 and 109.4 *±* 28.2 at P3 (paired two-sided *t*-tests, P1 versus P2: *t*(7) = 3.44, *P*_FDR_ = 0.0109; P1 versus P3: *t*(7) = 3.58, *P*_FDR_ = 0.0109). A similar attenuation was observed ipsilaterally (P1, 205.7 *±* 45.2; P2, 79.5 *±* 10.6; P3, 75.5 *±* 6.6; P1 versus P2: *t*(7) = 2.53, *P*_FDR_ = 0.0392; P1 versus P3: *t*(7) = 2.70, *P*_FDR_ = 0.0392; *n* = 8 rats; Fig. 2a,b).

**Fig. 2.**
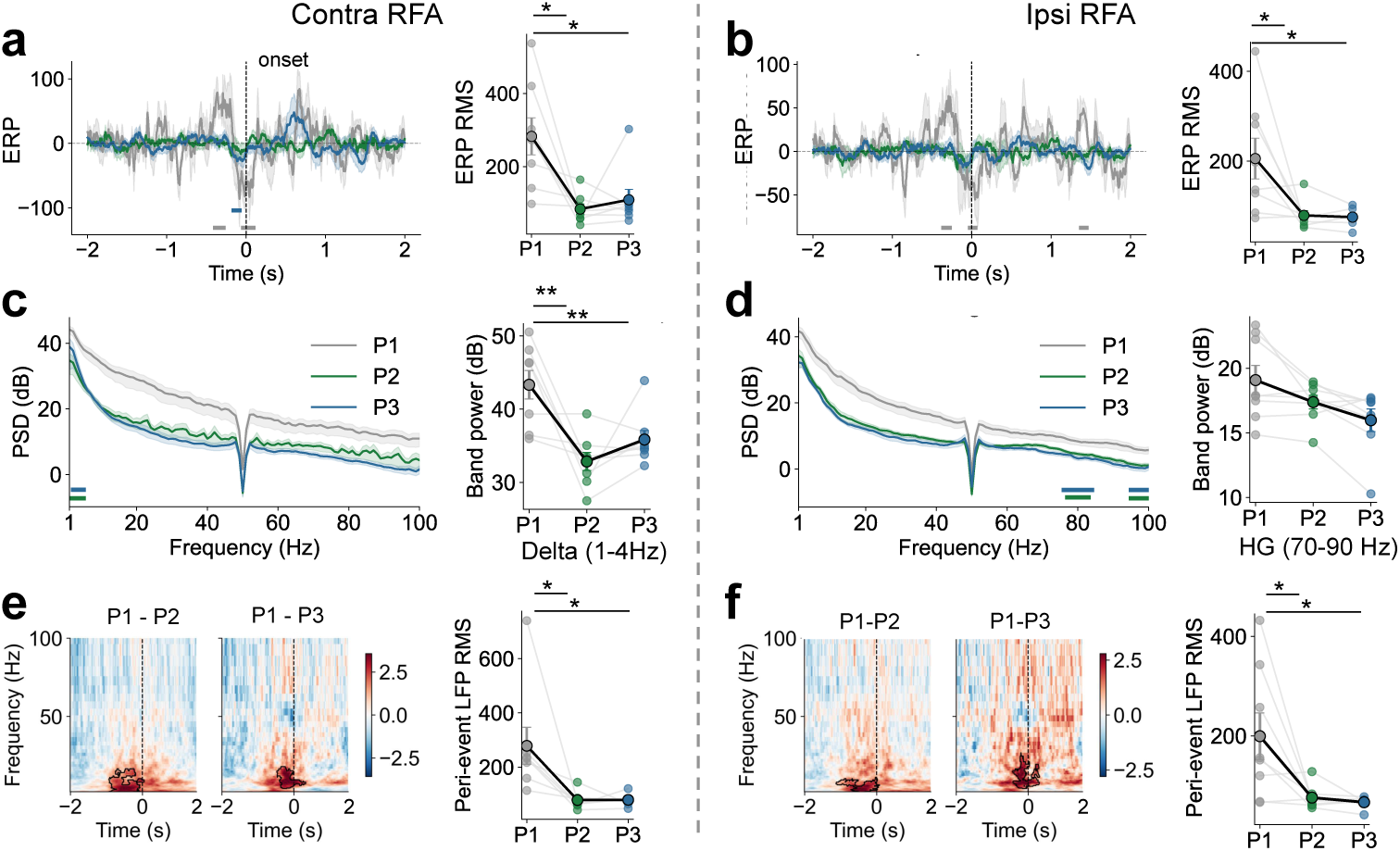
SPR-related LFP activity in bilateral RFA is attenuated across motor learning. a,b, Retrieval-aligned event-related potentials (ERPs) in contralateral (a) and ipsilateral (b) RFA across learning phases P1–P3. Time 0 denotes pellet retrieval onset; shaded regions indicate s.e.m. Right, LFP root-mean-square (RMS) amplitude quantified over the full analysis window. c,d, Power spectral density (PSD) in contralateral (c) and ipsilateral (d) RFA across learning phases. Horizontal bars indicate significant frequency clusters identified by trial-level cluster-based permutation tests. Right, rat-level delta-band power (1–4 Hz) in contralateral RFA (c) and high-gamma power (70–90 Hz) in ipsilateral RFA (d). e,f, Time–frequency differences in baseline-normalized LFP power between P1 and P2 and between P1 and P3 in contralateral (e) and ipsilateral (f) RFA. Power was normalized to the *−*3 to *−*2 s baseline and expressed in decibels. Black contours indicate significant time–frequency clusters identified by trial-level cluster-based permutation tests. Right, pre-event LFP RMS quantified over *−*2 to 0.3 s. For rat-level comparisons, lines connect measurements from the same animal and black symbols indicate mean *±* s.e.m. (*n* = 8 rats). Rat-level comparisons were performed using two-sided paired *t*-tests with false-discovery-rate (FDR) correction. *P <* 0.05 was considered significant.

We next examined the spectral organization of this attenuation. Power spectral density (PSD) decreased across learning in both hemispheres, but with distinct spectral profiles (Fig. 2c,d). In contralateral RFA, trial-level cluster-based permutation tests identified lower power at P2 than P1 over 1–5 Hz (*P* = 0.0154) and at P3 than P1 over 2–5 Hz (*P* = 0.0150). Rat-level delta-band power (1–4 Hz) confirmed this reduction (P1 versus P2: *t*(7) = 3.68, *P*_FDR_ = 0.0079; P1 versus P3: *t*(7) = 3.71, *P*_FDR_ = 0.0079). In ipsilateral RFA, significant trial-level differences were concentrated at higher frequencies, with P1–P2 clusters at 77–83 Hz (*P* = 0.0254) and 95–100 Hz (*P* = 0.0310), and P1–P3 clusters at 76–84 Hz (*P* = 0.0086) and 95–100 Hz (*P* = 0.0186). Rat-level high-gamma power (70–90 Hz) showed a corresponding decrease that did not survive FDR correction (P1 versus P2, *P*_FDR_ = 0.0856; P1 versus P3, *P*_FDR_ = 0.0792).

Time–frequency analysis further showed that the dominant learning-related differences were concentrated around pellet retrieval, with substantial effects emerging before retrieval onset (Fig. 2e,f). In contralateral RFA, power was lower at P2 than P1 over 2.0–15.9 Hz from −0.92 to 0.05 s (*P* = 0.0040) and at P3 than P1 over 4.1–17.3 Hz from −0.47 to 0.32 s (*P* = 0.0310). Ipsilateral RFA showed a similar reduction over 2.0–10.7 Hz from −1.12 to 0.08 s for P1 versus P2 (*P* = 0.0050) and 4.1–25.7 Hz from −0.43 to 0.31 s for P1 versus P3 (*P* = 0.0060). Consistent with this temporal localization, pre-event LFP RMS quantified over −2 to 0.3 s was reduced at P2 and P3 in both contralateral RFA (P1 versus P2: *t*(7) = 2.83, *P*_FDR_ = 0.0254; P1 versus P3: *t*(7) = 2.89, *P*_FDR_ = 0.0254) and ipsilateral RFA (P1 versus P2: *t*(7) = 2.52, *P*_FDR_ = 0.0400; P1 versus P3: *t*(7) = 2.87, *P*_FDR_ = 0.0400).

Because retrieval success differed across learning phases, we asked whether these LFP differences could reflect the changing proportion of successful and failed trials. We therefore repeated the analyses using successful retrieval trials only (Supplementary Fig. S1). The principal attenuation pattern was preserved. Whole-window LFP RMS remained reduced at P2 and P3 in contralateral RFA (P1 versus P2: *t*(7) = 3.45, *P*_FDR_ = 0.0156; P1 versus P3: *t*(7) = 3.18, *P*_FDR_ = 0.0156) and ipsilateral RFA (P1 versus P2: *t*(7) = 2.53, *P*_FDR_ = 0.0391; P1 versus P3: *t*(7) = 2.57, *P*_FDR_ = 0.0391). Contralateral low-frequency attenuation was likewise retained, with significant PSD clusters at 1–5 Hz for P1 versus P2 (*P* = 0.0102) and 3–5 Hz for P1 versus P3 (*P* = 0.0234), together with lower rat-level delta power at both later phases (*P*_FDR_ = 0.0099 for both comparisons). The ipsilateral high-gamma band showed the same numerical reduction but did not reach significance after FDR correction. Time–frequency differences nevertheless remained robust bilaterally: significant P1–P2 and P1–P3 clusters extended across low frequencies around retrieval in both contralateral and ipsilateral RFA, and pre-retrieval LFP RMS remained lower at both later phases contralaterally (*P*_FDR_ = 0.0215 for both comparisons) and ipsilaterally (*P*_FDR_ = 0.0388 for both comparisons). Thus, the attenuation of retrieval-related RFA activity persisted when behavioral outcome was held constant, indicating that it was not explained simply by the increasing proportion of successful trials across learning.

Together, these analyses show that improved SPR was accompanied by reduced, rather than progressively amplified, retrieval-related RFA recruitment. The effect was expressed bilaterally across complementary LFP measures, was particularly prominent around the pre-retrieval period, and remained evident when analyses were restricted to successful trials. This raised a key question for the underlying neuronal activity: did reduced population-level recruitment reflect diminishing premotor engagement, or did behaviorally relevant premotor dynamics persist as skill developed?

### 2.3 SPR-related firing suppression persists across learning and predicts successful retrieval

To determine whether the attenuation of retrieval-related RFA LFP activity reflected a loss of premotor engagement or a reorganization of the underlying neuronal population response, we next examined single-unit firing across learning. Population heat maps and unit-averaged firing profiles revealed a strikingly consistent pattern in both contralateral and ipsilateral RFA: retrieval-related activity was dominated by firing-rate suppression rather than excitation, and this suppressive motif remained evident throughout all three learning phases (Fig. 3a–b). Relative to baseline, significant firing suppression was present in every phase in both hemispheres. In contralateral RFA, suppression extended from 0.38 to 2.00 s in P1, from 2.00 to 2.00 s in P2, and from 2.00 to 1.48 s in P3 (all cluster *p* = 0.0002). In ipsilateral RFA, significant suppression was likewise observed from 0.94 to 2.00 s in P1, from 2.00 to 2.00 s in P2, and from 1.80 to 1.48 s in P3 (all cluster *p* = 0.0002). These results indicate that retrieval-related firing suppression is a stable physiological signature of bilateral RFA activity during skilled pellet retrieval.

**Fig. 3.**
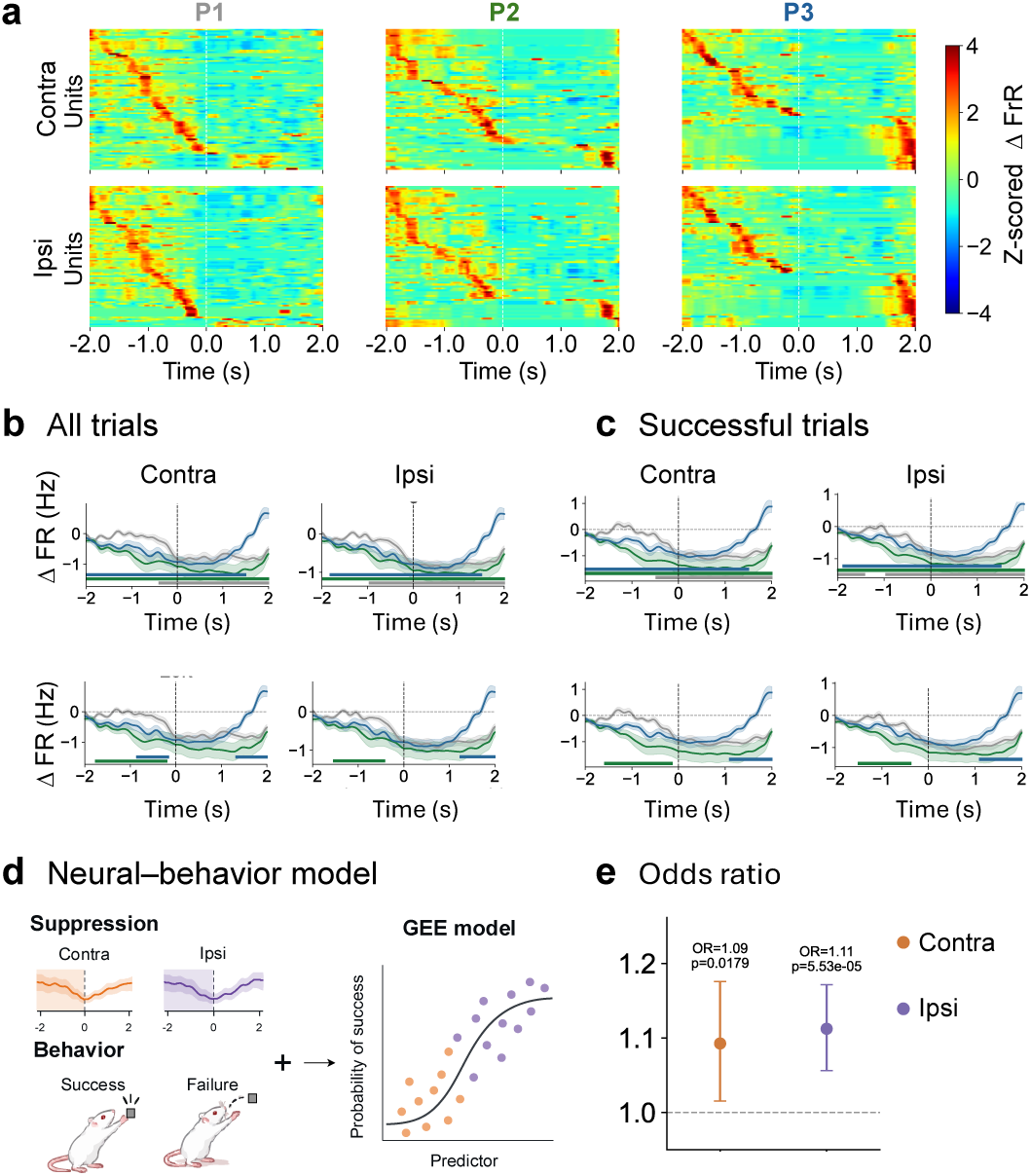
SPR-related firing suppression persists across motor learning and predicts successful retrieval. a, Population firing-rate activity in contralateral (upper) and ipsilateral (lower) RFA across learning phases P1–P3. Each row represents one unit and shows the baseline-corrected firing-rate change (Δ firing rate), z-scored within unit. Units were independently ordered within each phase according to the timing of their peak firing rate. Time 0 denotes pellet retrieval onset. b, Population-averaged Δ firing rate for all trials in contralateral (left) and ipsilateral (right) RFA. Curves show mean *±* s.e.m. across units for P1 (gray), P2 (green) and P3 (blue). Upper plots show significant periods of firing-rate suppression relative to baseline for each phase, identified by unit-level cluster-based permutation tests. Suppression was significant in all phases bilaterally (cluster *P* = 0.0002). Lower plots show learning-related differences between phases. Compared with P1, P2 exhibited stronger pre-retrieval suppression in contralateral RFA (*−*1.74 to *−*0.22 s, *P* = 0.019) and ipsilateral RFA (*−*1.52 to *−*0.44 s, *P* = 0.038). P3 differed from P1 primarily during the later retrieval-related period, with weaker suppression in contralateral RFA (1.34 to 2.00 s, *P* = 0.0166) and ipsilateral RFA (1.26 to 2.00 s, *P* = 0.0208); an additional contralateral cluster occurred from *−*0.84 to *−*0.18 s (*P* = 0.0464). c, The same analyses restricted to successful retrieval trials. Robust firing suppression remained evident across learning phases. Compared with P1, P2 again showed stronger pre-retrieval suppression in contralateral RFA (*−*1.56 to *−*0.16 s, *P* = 0.0214) and ipsilateral RFA (*−*1.48 to *−*0.40 s, *P* = 0.0386), whereas P3 showed weaker late suppression than P1 in contralateral RFA (1.12 to 2.00 s, *P* = 0.0060) and ipsilateral RFA (1.12 to 2.00 s, *P* = 0.0132). Horizontal colored bars denote significant temporal clusters. Analyses included 92 contralateral and 112 ipsilateral units. d, Schematic of the trial-level neural–behavior analysis relating pre-retrieval firing suppression to behavioral outcome. Suppression was quantified from baseline-corrected firing activity during the *−*2 to 0 s interval, with larger positive values indicating stronger firing-rate reduction, and entered into a generalized estimating equation (GEE) logistic model of retrieval success. e, Odds ratios (ORs) from the GEE models, expressed per 1-s.d. increase in pre-retrieval suppression. Stronger suppression was associated with higher odds of successful retrieval in both contralateral (OR = 1.09, *P* = 0.0179) and ipsilateral (OR = 1.11, *P* = 5.53 *×* 10*^−^*^5^) RFA. Points indicate estimated ORs and error bars indicate 95% confidence intervals; the dashed line denotes OR = 1.

Although suppression persisted across learning, its temporal expression was not identical across phases. Importantly, baseline firing rates did not differ significantly between phases in either hemisphere for all-trial analyses, indicating that the observed effects were not explained by nonspecific changes in spontaneous activity. Compared with P1, P2 exhibited stronger pre-retrieval suppression in both contralateral RFA (1.74 to 0.22 s, cluster *p* = 0.019) and ipsilateral RFA (1.52 to 0.44 s, cluster *p* = 0.038; Fig. 3b). By contrast, P3 differed from P1 mainly during the later retrieval-related period, with weaker suppression in contralateral RFA from 1.34 to 2.00 s (cluster *p* = 0.0166) and in ipsilateral RFA from 1.26 to 2.00 s (cluster *p* = 0.0208). An additional smaller contralateral difference was present before retrieval onset (0.84 to 0.18 s, cluster *p* = 0.0464). Thus, learning did not abolish suppressive firing in RFA, but was accompanied by phase-dependent adjustments in when that suppression was expressed.

Because trial outcome changed substantially across learning, we next asked whether this firing pattern would remain if the analysis was restricted to successful retrievals only. The successful-trial analysis yielded a highly similar overall picture (Fig. 3c). Significant suppression relative to baseline again remained evident across all phases in both hemispheres, with broad suppressive clusters spanning the retrieval period. Moreover, the key phase differences were recapitulated. P2 continued to show stronger pre-retrieval suppression than P1 in contralateral RFA (1.56 to 0.16 s, cluster *p* = 0.0214) and ipsilateral RFA (1.48 to 0.40 s, cluster *p* = 0.0386). P3 again showed weaker later suppression than P1 in contralateral RFA (1.12 to 2.00 s, cluster *p* = 0.0060) and ipsilateral RFA (1.12 to 2.00 s, cluster *p* = 0.0132). Together, these findings indicate that the persistence of retrieval-related firing suppression cannot be attributed simply to differences in the proportion of successful and failed trials across learning.

We then tested whether this suppressive activity was behaviorally relevant on a trial-by-trial basis. For each trial, we quantified pre-retrieval suppression over the 2 to 0 s interval and related it to retrieval outcome using a logistic model fitted with generalized estimating equations (GEE; Fig. 3d,e). In both hemispheres, stronger pre-retrieval suppression was associated with a greater probability of successful retrieval. In contralateral RFA, a 1-s.d. increase in suppression was associated with higher odds of successful retrieval (odds ratio (OR), 1.09; 95% confidence interval (CI), 1.015– 1.176; *P* = 0.0179). In ipsilateral RFA, a 1-s.d. increase in suppression was likewise associated with higher odds of successful retrieval (OR, 1.11; 95% CI, 1.056–1.171; *P* = 5.53 10*^−^*^5^). Thus, although large-scale retrieval-related LFP activity became attenuated as behavior improved, the underlying suppressive firing motif remained robust and continued to predict successful performance.

Overall, these results show that the main single-unit signature of bilateral RFA during pellet retrieval is not lost across learning. Instead, retrieval-related firing suppression persists across learning, remains evident when successful trials are considered alone, and retains clear behavioral relevance at the trial level. This finding argues against a simple interpretation in which reduced population-level RFA activity reflects diminished involvement. Rather, it suggests that a functionally meaningful suppressive control signal is maintained even as overt population recruitment becomes attenuated.

### 2.4 SPR-related suppression is preferentially expressed by broad-spiking RFA units

Having established that SPR-related firing suppression persisted across learning and was associated with successful retrieval, we next asked whether this suppressive motif reflected a uniform reduction in RFA firing or was expressed by a selective neuronal population. Representative longitudinal recordings illustrated units with pronounced suppression around pellet retrieval, while the expression of this response differed across learning phases (Fig. 4a). We therefore identified units exhibiting significant decreases in firing relative to baseline and quantified their prevalence within each hemisphere.

**Fig. 4.**
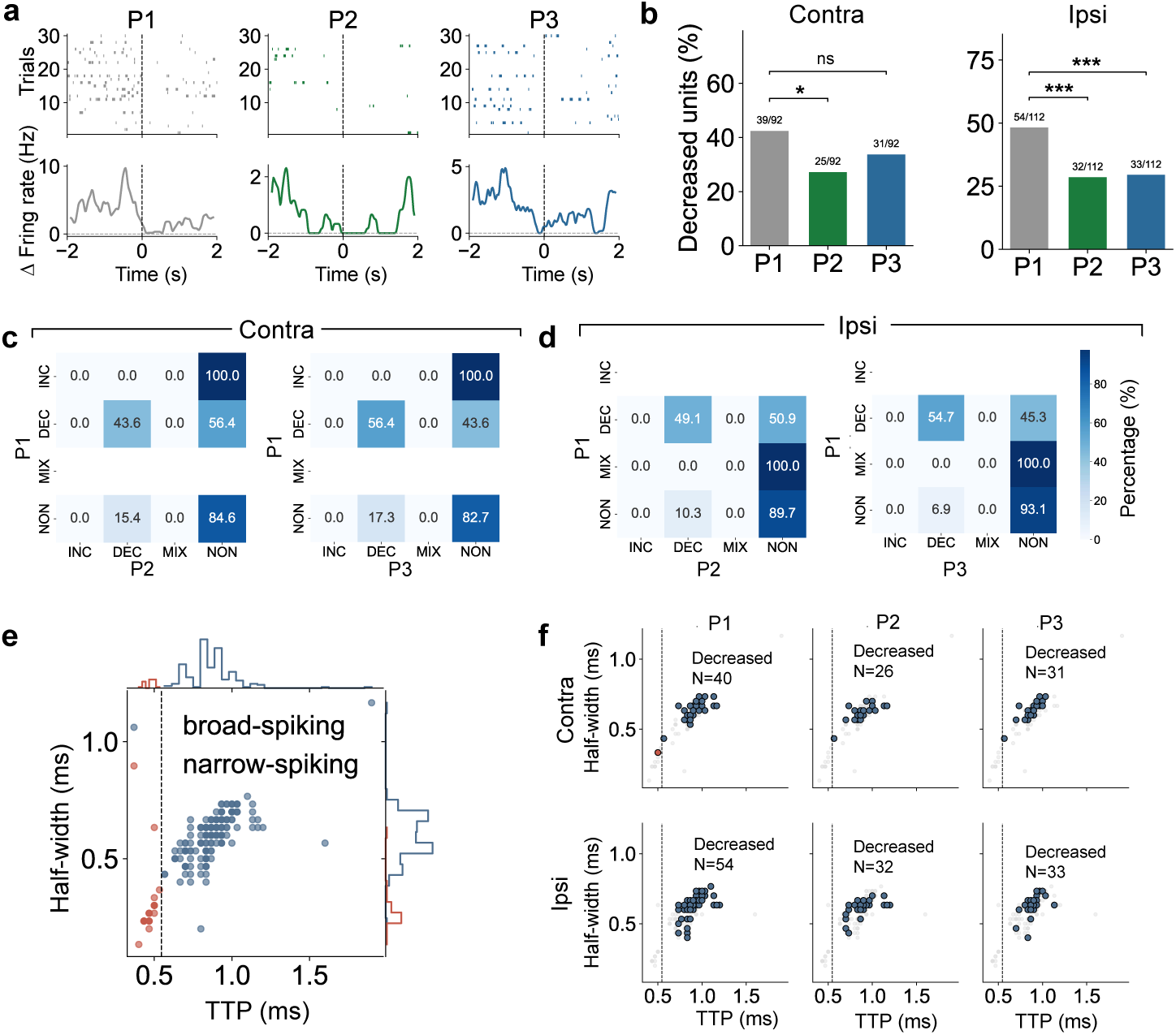
Learning restricts and reorganizes the neuronal population recruited into retrieval-related firing suppression. **a**, Representative firing activity of a decreased unit across learning phases P1–P3. Upper panels show spike rasters across trials, and lower panels show the corresponding baseline-corrected firing-rate change (Δ firing rate). Time 0 denotes pellet retrieval onset. **b**, Proportion of units classified as decreased in contralateral and ipsilateral RFA across learning phases. In contralateral RFA, decreased units comprised 39/92 (42.4%), 25/92 (27.2%) and 31/92 (33.7%) units in P1, P2 and P3, respectively. The proportion was reduced from P1 to P2 (*P*_FDR_ = 0.0323), whereas the difference between P1 and P3 was not significant (*P*_FDR_ = 0.169). In ipsilateral RFA, decreased units comprised 54/112 (48.2%), 32/112 (28.6%) and 33/112 (29.5%) units in P1, P2 and P3, respectively, with significant reductions from P1 to both P2 (*P*_FDR_ = 1.95 *×* 10*^−^*^4^) and P3 (*P*_FDR_ = 1.95 *×* 10*^−^*^4^). Paired proportions were compared using two-sided exact McNemar tests followed by FDR correction. ns, not significant; \**P <* 0.05; \*\*\**P <* 0.001. **c,d**, Transition matrices showing changes in firing-response classification from P1 to P2 and from P1 to P3 for the same units in contralateral **c** and ipsilateral **d** RFA. Units were classified as increased (INC), decreased (DEC), mixed (MIX) or non-modulated (NON) according to their SPR-related firing-rate responses. Rows indicate the P1 classification and columns indicate the classification in P2 or P3; values show the percentage of units within each P1 category transitioning to each later-phase category.**e**, Waveform-based classification of recorded units. Gaussian mixture modelling of trough-to-peak time (TTP) identified a cutoff of 0.546 ms separating narrow- and broad-spiking units; waveform half-width is shown as a complementary waveform feature. Marginal distributions show the corresponding TTP and half-width distributions. **f**, Waveform-feature distributions of suppression-expressing units across learning phases and hemispheres. In contralateral RFA, 38 of 39 suppression-expressing units at P1 and all suppression-expressing units at P2 (25 of 25) and P3 (31 of 31) were broad-spiking. In ipsilateral RFA, all suppression-expressing units were broad-spiking at P1 (54 of 54), P2 (32 of 32) and P3 (33 of 33). Suppression was more prevalent among broad-spiking than narrow-spiking units at every phase in both hemispheres (two-sided Fisher’s exact tests with FDR correction).

Only a subset of recorded units expressed significant SPR-related suppression (Fig. 4b). In contralateral RFA, decreased units comprised 39 of 92 units (42.4%) at P1, 25 of 92 (27.2%) at P2 and 31 of 92 (33.7%) at P3. The proportion was lower at P2 than P1 (*P*_FDR_ = 0.0323, exact McNemar test), whereas the P1–P3 difference was not significant (*P*_FDR_ = 0.1686). In ipsilateral RFA, the corresponding proportions were 54 of 112 units (48.2%) at P1, 32 of 112 (28.6%) at P2 and 33 of 112 (29.5%) at P3, with significant reductions from P1 to both P2 and P3 (*P*_FDR_ = 1.95 10*^−^*^4^ for both comparisons). Thus, the behaviorally relevant suppressive motif identified in Fig. 3 was expressed by a restricted fraction of the RFA population, with fewer neurons exhibiting suppression at the later learning phases.

We next examined whether this suppression was carried by a fixed neuronal ensemble across learning. Same-unit transition analyses revealed substantial turnover in suppression status (Fig. 4c,d). In contralateral RFA, 43.6% of units classified as decreased at P1 remained decreased at P2 and 56.4% remained decreased at P3, whereas the remaining P1 decreased units became non-modulated. Conversely, 15.4% and 17.3% of units that were non-modulated at P1 expressed suppression at P2 and P3, respectively. A similar pattern was observed ipsilaterally: 49.1% of P1 decreased units remained decreased at P2 and 54.7% at P3, while 10.3% and 6.9% of initially non-modulated units became decreased at the corresponding later phases. Thus, SPR-related suppression was not confined to a fixed set of neurons; rather, membership in the suppression-expressing population remained selective across learning.

To determine whether SPR-related suppression was preferentially expressed by a waveform-defined neuronal population, we classified units as broad- or narrow-spiking using Gaussian mixture modelling of trough-to-peak time (TTP), with waveform half-width providing a complementary waveform feature (Fig. 4E). Broad-spiking units constituted 81 of 92 units (88.0%) in contralateral RFA and 99 of 112 units (88.4%) in ipsilateral RFA. Suppression-expressing units were almost exclusively broad-spiking. In contralateral RFA, 38 of 39 suppression-expressing units (97.4%) at P1 and all such units at P2 (25 of 25) and P3 (31 of 31) were broad-spiking. In ipsilateral RFA, all suppression-expressing units were broad-spiking at each phase (P1, 54 of 54; P2, 32 of 32; P3, 33 of 33; Fig. 4F). At the unit level, suppression was more prevalent among broad-spiking than narrow-spiking units at every phase in both hemispheres (two-sided Fisher’s exact tests; contralateral P1–P3, all *P*_FDR_ = 0.0318; ipsilateral P1, *P*_FDR_ = 3.76 10*^−^*^4^; P2, *P*_FDR_ = 0.0182; P3, *P*_FDR_ = 0.0146). Thus, SPR-related suppression was preferentially associated with the broad-spiking extracellular population.

Together, these findings refine the population-level picture emerging from the preceding analyses. Although SPR-related suppression remained a robust and behaviorally relevant feature of RFA activity, it was implemented by a selective neuronal subset whose recruitment was more restricted at later learning phases and whose membership was not fixed across learning. This raised the next question of whether the overall population neural state, beyond the activity of individual suppression-expressing neurons, showed a corresponding learning-stage-dependent relationship with successful retrieval.

### 2.5 Successful retrieval is most strongly coupled to premotor population neural state engagement early in learning

The selective recruitment of suppression-expressing neurons suggested that learning might also alter how strongly the broader RFA population is engaged during individual retrieval attempts. We therefore examined trial-resolved population activity in low-dimensional state space and quantified its departure from a pre-retrieval reference state using multidimensional distance (MDD; Fig. 5a–d). In both contralateral and ipsilateral RFA, population trajectories departed from the reference state before pellet retrieval and remained displaced around retrieval onset, indicating coordinated population-level engagement during SPR.

**Fig. 5.**
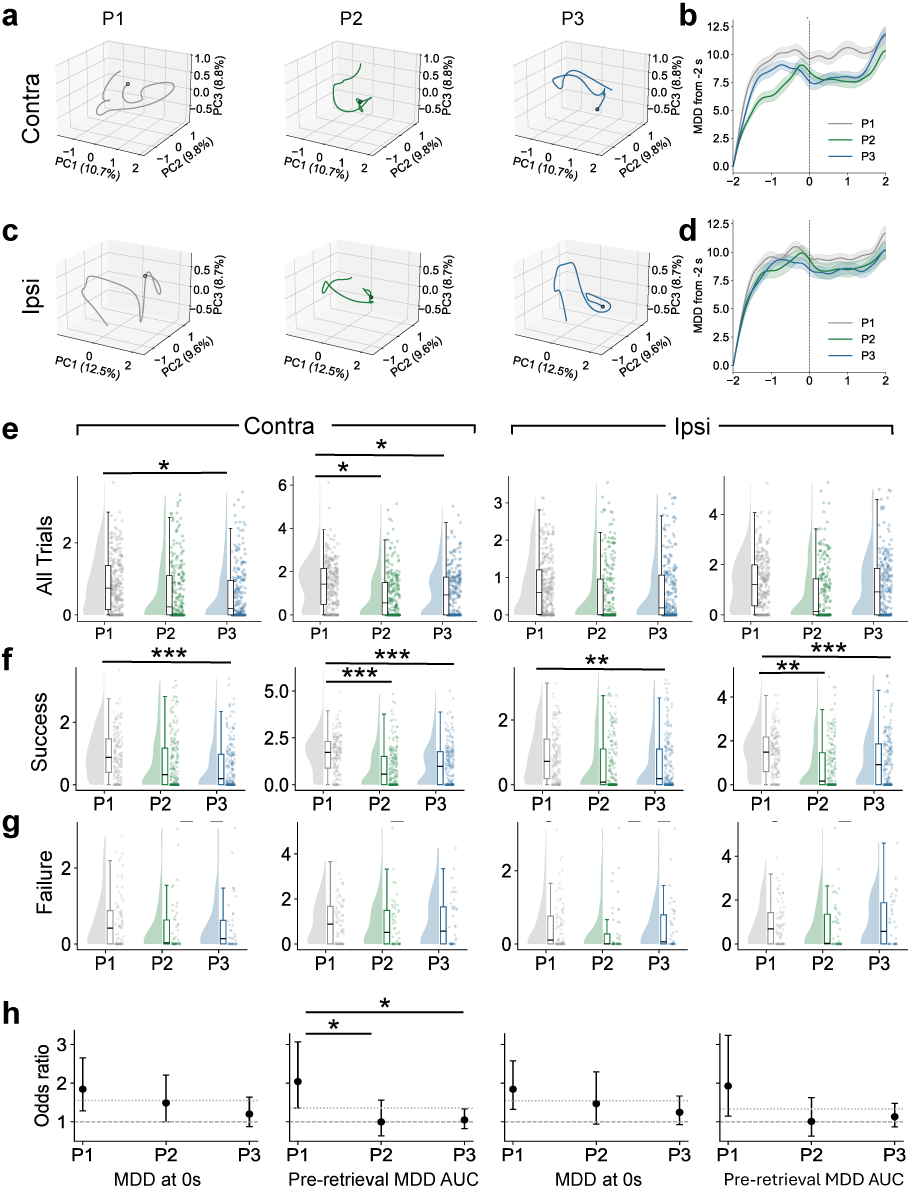
Premotor population neural state engagement is most strongly associated with successful retrieval early in learning. a,C, Population trajectories in the first three principal components (PCs) for contralateral (a) and ipsilateral (c) RFA across learning phases P1–P3. Trajectories were constructed from retrieval-aligned population firing activity over the *−*2 to 2 s interval; time 0 denotes pellet retrieval onset. b,d, Time-resolved multidimensional distance (MDD) of the population neural state from its state at *−*2 s in contralateral (b) and ipsilateral (d) RFA. Curves show mean *±* s.e.m. for P1 (gray), P2 (green) and P3 (blue). e, Trial-level distributions of MDD at retrieval onset (MDD at 0 s) and pre-retrieval MDD area under the curve (Pre-AUC; *−*2 to 0 s) across learning phases for all trials. Raincloud plots show individual trials, distributions and summary box plots. In contralateral RFA, MDD at 0 s was lower in P3 than P1 (*P*_FDR_ = 0.0385), and Pre-AUC was lower in both P2 and P3 than P1 (*P*_FDR_ = 0.0379 for both comparisons). No phase comparison survived FDR correction in ipsilateral RFA. f, The same analysis restricted to successful retrieval trials. Successful trials showed lower MDD at 0 s in P3 than P1 in both contralateral (*P*_FDR_ = 6.0 *×* 10*^−^*^6^) and ipsilateral (*P*_FDR_ = 0.00108) RFA. Pre-AUC was reduced from P1 to P2 and P3 in contralateral RFA (*P*_FDR_ = 2.54 *×* 10*^−^*^4^ and 7.0 *×* 10*^−^*^6^, respectively) and ipsilateral RFA (*P*_FDR_ = 0.00189 and 6.4 *×* 10*^−^*^5^, respectively). g, Corresponding distributions for failed retrieval trials; no phase comparison survived FDR correction. Phase effects in e–g were evaluated using generalized estimating equation (GEE) models accounting for repeated trials within rats, followed by FDR-corrected pair-wise contrasts. h, Phase-specific associations between population neural state metrics and retrieval success estimated using logistic GEE models. Points indicate odds ratios for successful retrieval per 1-s.d. increase in each metric and error bars indicate 95% confidence intervals; the dashed horizontal line denotes an odds ratio of 1. Both MDD at 0 s and Pre-AUC were positively associated with success during P1 in contralateral and ipsilateral RFA. The association between Pre-AUC and success varied significantly across learning phases in contralateral RFA (metric *×* phase interaction, *P* = 0.0202), with a stronger association in P1 than P2 (OR ratio = 2.05, *P*_FDR_ = 0.0203) or P3 (OR ratio = 1.95, *P*_FDR_ = 0.0203). Corresponding phase interactions for MDD at 0 s and for ipsilateral Pre-AUC were not significant. All analyses included *n* = 8 rats. \**P <* 0.05, \*\**P <* 0.01, \*\*\**P <* 0.001.

We first asked whether the magnitude of this engagement differed across learning. Two complementary measures were considered: MDD at retrieval onset and the area under the MDD trajectory during the pre-retrieval period (Pre-AUC; Fig. 5e). In contralateral RFA, MDD at 0 s showed a significant phase effect (Wald *χ*^2^(2) = 7.71, *p* = 0.0212), with lower engagement at P3 than P1 (*P*_FDR_ = 0.0385). Pre-AUC showed the same overall tendency: although the omnibus phase effect was marginal (*p* = 0.0517), pairwise contrasts revealed lower sustained engagement at both P2 and P3 relative to P1 (*P*_FDR_ = 0.0379 for both comparisons). Corresponding phase effects did not reach significance in ipsilateral RFA. Thus, the attenuation observed in the LFP analyses was also evident in the population neural state dynamics, particularly in contralateral RFA.

We next separated successful and failed retrievals to determine whether this learning-stage dependence was linked to behavioral outcome. For successful trials, both MDD measures varied strongly across phases in both hemispheres (contralateral: MDD at 0 s, Wald *χ*^2^(2) = 22.07, *p* = 1.6 10*^−^*^5^; Pre-AUC, *χ*^2^(2) = 22.77, *p* = 1.1 10*^−^*^5^; ipsilateral: MDD at 0 s, *χ*^2^(2) = 12.27, *p* = 0.00216; Pre-AUC, *χ*^2^(2) = 22.35, *p* = 1.4 10*^−^*^5^; Fig. 5f). Successful P1 trials showed greater sustained pre-retrieval engagement than successful trials at both P2 and P3 in contralateral RFA (*P*_FDR_ = 2.54 10*^−^*^4^ and 7 10*^−^*^6^, respectively) and ipsilateral RFA (*P*_FDR_ = 0.00189 and 6.4 10*^−^*^5^, respectively). MDD at 0 s was likewise higher at P1 than P3 bilaterally. In contrast, neither MDD at 0 s nor Pre-AUC differed significantly across phases for failed trials (Fig. 5g). The learning-related reduction in population neural state engagement was therefore expressed primarily in successful actions rather than uniformly across all retrieval attempts.

We then asked whether trial-to-trial variation in population neural state engagement predicted retrieval success. Across all phases, greater engagement was associated with higher odds of successful retrieval for both metrics and both hemispheres (Fig. 5h). For MDD at 0 s, For MDD at 0 s, the overall odds ratios (ORs) were 1.55 in contralateral RFA (*p* = 2.2 10*^−^*^5^) and 1.55 ipsilaterally (*p* = 2.2 10*^−^*^5^). Pre-AUC was similarly associated with success (contralateral, OR = 1.36, *p* = 0.00132; ipsilateral, OR = 1.33, *p* = 0.00684). This relationship was strongest during early learning: at P1, MDD at 0 s was associated with approximately 1.85-fold greater odds of success in both contralateral (OR = 1.85, *P*_FDR_ = 0.00294) and ipsilateral RFA (OR = 1.84, *P*_FDR_ = 9.94 *×* 10*^−^*^4^), while Pre-AUC was associated with a 2.04-fold increase contralaterally (*P*_FDR_ = 0.00190) and a 1.93-fold increase ipsilaterally (*P*_FDR_ = 0.0385).

The learning dependence of this behavioral coupling was clearest for sustained pre-retrieval engagement. In contralateral RFA, the relationship between Pre-AUC and retrieval success differed significantly across phases (interaction, Wald *χ*^2^(2) = 7.81, *p* = 0.0202), with the P1 association approximately twice as strong as that at P2 (OR ratio = 2.05, *P*_FDR_ = 0.0203) and P3 (OR ratio = 1.95, *P*_FDR_ = 0.0203). By contrast, phase interactions were not significant for MDD at 0 s in either hemisphere, and the corresponding Pre-AUC interaction did not reach significance ipsilaterally. Thus, the clearest learning-dependent shift concerned the behavioral relevance of sustained pre-retrieval population engagement, rather than the instantaneous population displacement at retrieval itself.

Together, these findings show that RFA population neural state engagement was most strongly coupled to successful retrieval early in learning. As performance improved, successful actions were accompanied by less sustained population recruitment, while the behaviorally relevant suppressive firing motif remained preserved. This pattern completes the progression across the preceding results, supporting a shift from broad premotor population engagement during early acquisition toward a more selective organization of premotor control as skill became established.

## Discussion

Practice transforms variable and unreliable actions into increasingly reliable skilled behavior [17, 19]. Yet how premotor cortex contributes to the control required for this reliability remains unclear. Does reliable skilled action require progressively stronger premotor recruitment, or can it emerge with less widespread cortical engagement? Here, we followed bilateral RFA activity while rats acquired the SPR task. As performance improved, broad SPR-related LFP activity was attenuated rather than amplified (Fig. 2). At the same time, single-unit activity remained dominated by SPR-related firing suppression, and stronger pre-retrieval suppression was associated with greater odds of successful retrieval (Fig. 3). Thus, improved skill was accompanied by attenuated broad premotor activity without a corresponding loss of behaviorally relevant premotor dynamics.

One straightforward interpretation of the reduced SPR-related activity is that RFA becomes progressively less involved once the task is learned. Several observations argue against this interpretation. First, the attenuation of LFP activity was preserved when analyses were restricted to successful trials (Supplementary Fig. S1), indicating that it could not be explained simply by the increasing proportion of successful attempts across learning. Second, robust SPR-related firing suppression remained present in both hemispheres throughout P1–P3 despite the reduction in broader population activity (Fig. 3). Third, this suppressive response retained behavioral relevance at the level of individual trials: stronger pre-retrieval suppression was associated with greater odds of successful retrieval even after accounting for learning phase. More broadly, motor-learning studies have shown that cortical participation need not increase uniformly with practice. In primary motor cortex, an initially expanded and variable movement-related population can be refined into a smaller population expressing more reproducible activity patterns as learning proceeds [7], and different cortical layers can follow distinct trajectories of ensemble reorganization [8]. Cortical involvement can also depend strongly on task demands and learning stage; for example, motor cortex is required for acquiring, but not subsequently executing, a learned non-dexterous motor sequence [18]. Together, these observations argue against equating reduced cortical activity with reduced functional involvement. Instead, RFA attenuation appears to reflect a reorganization of premotor participation as reliable behavior emerges.

SPR-related firing suppression provides a potential account of how RFA contributes to skilled action. Premotor activity is often considered in the context of action specification and selection, but selection need not be expressed solely through increased firing associated with a chosen movement. In primate dorsal premotor cortex, multiple potential reaching actions can be represented concurrently, and commitment to one action is accompanied by enhancement of the selected representation together with attenuation of the rejected alternative [20, 21]. Such observations establish a broader principle that reductions in premotor firing can participate in active control rather than simply reflect neuronal inactivity. Our SPR task differs importantly from explicit choice paradigms: the animals were not choosing between simultaneously instructed reach targets, and the suppressed activity observed here therefore cannot be identified with a specific rejected action. Nevertheless, a purely passive interpretation is difficult to reconcile with the behavioral relationship in our data, because trials with stronger pre-retrieval suppression were more likely to result in successful pellet retrieval (Fig. 3). One possibility is that SPR-related suppression reflects a restriction of premotor population activity as an ongoing action converges on an effective motor solution, reducing activity that is unnecessary or incompatible with successful performance. A reduction in extracellular firing does not identify the underlying circuit mechanism and should not be equated directly with increased synaptic inhibition. Rather, SPR-related suppression may represent a physiological signature of premotor constraint at the population level.

If suppression reflects a meaningful premotor operation, its persistence does not necessarily imply that the same neurons implement that operation throughout learning. Indeed, the neuronal expression of suppression was selective and dynamically organized. Only a subset of recorded RFA units showed significant SPR-related decreases in firing, and recruitment into this suppressive population was reduced most clearly from P1 to P2, with a sustained reduction in ipsilateral RFA at P3 (Fig. 4). Longitudinal tracking further showed substantial transitions between decreased and non-decreased response classes, indicating that persistence of the suppressive motif did not depend on a fixed neuronal ensemble. Thus, a functional pattern can be maintained while the neuronal population expressing it is reorganized. This distinction resembles a broader feature of motor learning in which stable or reproducible population-level representations can emerge despite substantial dynamism at the level of individual neurons [7]. Suppression-expressing units were almost exclusively broad-spiking across learning phases and hemispheres (Fig. 4). Together, these findings indicate that the suppressive motif was maintained by a selective but dynamically reorganized neuronal population. The data therefore point more strongly to a reorganization of the neuronal population expressing suppression than to a specific cellular mechanism.

The amount of population-level RFA engagement associated with successful action also depended on the stage at which the skill was expressed. Population trajectories departed from their pre-retrieval reference state in all phases, but successful trials showed substantially less population neural state engagement at later phases, whereas comparable phase-dependent differences were not evident for failed trials (Fig. 5). More importantly, trial-to-trial variation in population neural state was systematically associated with behavioral outcome. Early in learning, greater MDD at retrieval onset and greater sustained pre-retrieval MDD were associated with higher odds of successful retrieval. The learning-dependent difference was clearest for sustained pre-retrieval engagement: in contralateral RFA, the relationship between Pre-AUC and success was significantly stronger at P1 than at P2 or P3, whereas the corresponding phase dependence was not evident for the instantaneous MDD measured at retrieval onset. These findings suggest that early successful performance is particularly coupled to sustained recruitment of the premotor population before retrieval. As reliable performance emerges, the same behavioral outcome is achieved with smaller population neural state excursions. This reduction in broad engagement is notable because the SPR-related suppressive motif remained behaviorally relevant. The two observations together suggest that reliable skill does not arise through a uniform reduction of premotor involvement; rather, successful behavior becomes less strongly coupled to sustained population-wide engagement while a more specific control-related firing motif is retained.

This organization of RFA activity also informs how premotor cortex should be considered within the cortical forelimb network. RFA and CFA are strongly interacting forelimb representations, and recent simultaneous recording and perturbation experiments support a partial functional hierarchy between the two regions, with asymmetric inter-regional influence despite extensive sharing of firing patterns during reaching [15]. Our findings add a complementary dimension to this picture. When RFA activity was aligned to pellet retrieval onset and examined across skill acquisition, its dominant single-unit signature was SPR-related suppression rather than a uniformly excitatory movement-related response. This difference need not indicate a contradiction. Activity aligned to pellet retrieval onset samples a different portion of the behavioral sequence from activity aligned to movement initiation, and the expression of RFA activity is also likely to depend on task demands, learning stage, cortical sampling and the behavioral variable used for alignment. More importantly, recent work showing that functional hierarchy between RFA and CFA is not necessarily apparent from their firing patterns alone cautions against interpreting similarity or dissimilarity in firing profiles as a direct readout of hierarchical influence [15]. RFA may therefore be better understood not as a simple upstream replica of CFA, but as a cortical node whose population dynamics participate in control operations that depend on the demands of skilled action. The bilateral expression of SPR-related suppression and its association with successful retrieval suggest that this premotor contribution extends beyond the hemisphere contralateral to the reaching forelimb. The clearer learning dependence of sustained population neural state engagement in contralateral RFA further points to an asymmetric organization of this bilateral contribution.

The present findings support a model in which successful performance does not depend on progressively stronger premotor recruitment. Early in skill acquisition, successful actions are accompanied by broad and sustained RFA population engagement. As reliable performance emerges, broad SPR-related recruitment is attenuated and the neuronal population expressing suppression becomes more selective and reorganized, yet pre-retrieval suppression remains associated with successful behavior. We therefore propose that skilled action is supported by a transition from broadly recruited premotor control during early acquisition toward a more selective implementation of premotor constraint once reliable performance is established. This interpretation remains a population-level model rather than a direct circuit mechanism. The association between suppression and successful retrieval does not establish that suppression is causally required for performance, and extracellular waveform features do not identify the cell types generating this response. Moreover, because neural activity was aligned to pellet retrieval onset rather than movement onset, the pre-retrieval activity described here should not be interpreted simply as movement-preparatory activity. Combining longitudinal recordings with detailed movement kinematics, simultaneous measurements across the RFA–CFA network and temporally targeted perturbations will be important for determining which components of skilled action are constrained by RFA and how this control is implemented across the broader motor system. Motor learning therefore refines the neuronal and temporal organization of premotor cortical constraint as skilled behavior becomes reliable.

## Methods

### Animals and experimental design

Twenty-four adult male Sprague–Dawley rats (10 weeks old, 280–330 g; Zhuhai Bestest Biotechnology Company) were used in this study. Of these, eight rats that exhibited a consistent right-forelimb preference, achieved behavioral proficiency, and retained high-quality bilateral electrophysiological recordings across all three longitudinal learning phases (P1–P3) were included in the final behavioral and neural analyses (n = 8). Animals were housed in a specific pathogen-free facility and maintained at 21 2 *^◦^*C under a 12-h light/12-h dark cycle. Water was available ad libitum. Food intake was moderately restricted during behavioral training and recording periods to maintain motivation for the SPR task, while body weight was monitored daily and maintained above 90% of the pre-restriction baseline.

All animal procedures were approved by the Animal Experimental Ethics Committee of the Third Affiliated Hospital of Sun Yat-sen University and Guangdong Institute of Advanced Biomaterial and Medical Device (License No: IBMD-ES-25002) and were performed in accordance with institutional guidelines for animal care and use.

The longitudinal experimental design is shown in Fig. 1a. Recording electrodes were implanted bilaterally in the rostral forelimb area (RFA) at Week 0. Neural activity and behavioral performance were subsequently assessed during three learning phases: P1 at Week 1, P2 at Week 3 and P3 at Week 5. Local field potentials (LFPs) and single-unit spiking activity were recorded simultaneously during performance of the SPR task at each phase.

### Behavioral training

Rats were trained to perform the SPR task, a widely used paradigm for assessing skilled forelimb behavior [22], using a custom-built automated training system. The apparatus consisted of a behavioral chamber (40 cm 30 cm 25 cm) with a vertical slit (8 cm high, 1.2 cm wide) in the front wall. An automated feeding device was positioned 0.5 cm in front of the slit and controlled by a programmable microcontroller unit. Food pellets (45 mg Dustless Precision Pellets, Bio-Serv, product no. F0023) were delivered to a fixed target position by a robotic arm. Sensors positioned within the feeding device continuously monitored pellet displacement and generated digital event signals for behavioral synchronization. To evaluate the kinematics of the reaching and grasping behavior, we tracked the forelimb and paw trajectories using DeepLabCut, a deep-learning-based markerless pose estimation software [23].

At the beginning of training, forelimb preference was evaluated across repeated spontaneous reaching attempts (*>* 70% of reaching attempts made with the right paw). Animals displaying ambidexterity or left-paw preference were excluded from chronic implantation to maintain anatomical consistency across contralateral and ipsilateral hemispheres.

At the start of each trial, a pellet was positioned at the target location and an auditory cue was presented. The front baffle then opened, allowing the animal to reach through the slit and attempt pellet retrieval. Displacement of the pellet from the target position was detected by the feeding-device sensors and generated a transistor– transistor logic (TTL) pulse. This signal simultaneously triggered closure of the baffle, initiated resetting of the feeding system and was transmitted to the electrophysiological acquisition system. The TTL pulse was operationally defined as pellet retrieval onset and served as time 0 for all SPR-related neural analyses (Fig. 1d). Each trial was classified as successful or failed according to whether the animal successfully retrieved the pellet.

Animals continued to perform the SPR task throughout the longitudinal recording period. During daily behavioral sessions, each rat was presented with up to 30 pellets. Additional food was provided after each session to maintain body weight above 90% of baseline. Water remained available ad libitum.

### Electrode implantation

Following behavioral training, rats underwent chronic bilateral implantation of micro-filament electrode arrays targeting the RFA. Animals were anesthetized with isoflurane (4% induction and 2% maintenance concentrations) and positioned in a stereotaxic apparatus. Body temperature was maintained throughout surgery using a feedback-controlled heating pad. After aseptic preparation, a midline scalp incision was made and the skull was exposed. A craniotomy was created over each RFA. The centers of the left and right arrays were targeted to the RFA (AP = +2.5 mm, ML = ±2.5 mm relative to bregma), based on coordinates established in previous intracortical microstimulation mapping studies[11][24]. Additional skull openings were made for ground and reference screws.

The recording system comprised 16 microfilament electrodes distributed bilaterally (KD-BC, Suzhou), with one 2×4 array positioned in each hemisphere. Electrodes were fabricated from 35-*µ*m nickel–titanium wire and had impedances of approximately 100–200 kΩ before implantation. After opening the dura, arrays were lowered slowly into the cortex to a depth of 1550 ± 100 *µ*m from the cortical surface, avoiding visible surface vasculature. The arrays and connector were secured to the skull using dental cement.

### Electrophysiological recordings

Extracellular neural signals were recorded using the CereCube system (NeuroXess., Shanghai, China). Broadband extracellular activity was acquired from all implanted channels at a sampling rate of 30 kHz. Recordings were performed while animals freely executed the SPR task inside a copper-mesh Faraday cage, and the acquisition system was electrically grounded to minimize environmental interference.

Behavioral TTL pulses generated by pellet displacement were recorded synchronously with the neural signals and used to align electrophysiological activity to pellet retrieval onset. Neural activity was recorded bilaterally from RFA during P1, P2 and P3 using the same chronically implanted arrays.

### Spike sorting and single-unit quality control

Single-unit activity was isolated from the high-frequency component of the extracellular recordings (300–3,000 Hz). To enable longitudinal comparison of neuronal activity across learning, recordings obtained from the same electrode channel at P1, P2 and P3 were concatenated before spike sorting and processed using a common clustering solution. Spike sorting was performed with MountainSort4 implemented through the SpikeInterface framework [25, 26]. Following sorting, spike times were reassigned to their corresponding recording phases for phase-specific and longitudinal analyses.

Unit quality was evaluated using spike count and refractory-period violations [27]. Putative single units were retained if they contained more than 100 detected spikes and fewer than 1% of inter-spike intervals shorter than 2 ms. Waveforms and spike trains were additionally inspected to exclude poorly isolated clusters and obvious recording or sorting artifacts. Summary quality-control measures for retained units are shown in Supplementary Fig. S2.

### Local field potential preprocessing

LFP signals were sampled at 1,000 Hz and processed offline using MNE-Python [28]. Power-line interference was attenuated using a 50-Hz notch filter with a 2-Hz notch width, and signals were subsequently low-pass filtered at 200 Hz. LFP signals were aligned to behavioral events using TTL pulses generated by pellet displacement, which defined pellet retrieval onset (time 0). Continuous LFP recordings were segmented from 3 to +2 s relative to retrieval onset. No baseline correction was applied during epoch extraction; baseline normalization was instead performed separately according to the requirements of the event-related, spectral and time–frequency analyses. Signals from the contralateral and ipsilateral hemispheres were analyzed separately in subsequent analyses.

### Neural data analysis

#### Event-aligned LFP analysis

Event-aligned LFP activity was analyzed separately in the contralateral and ipsilateral RFA. Within each trial, signals were averaged across channels in the corresponding hemisphere, and trial-wise traces were subsequently averaged within each animal and learning phase to obtain animal-level event-aligned responses.

LFP responses were referenced to a pre-retrieval baseline (3 to 2 s relative to pellet retrieval onset). For each animal, hemisphere and learning phase, the mean baseline amplitude across trials was subtracted from the corresponding event-aligned response. Analyses focused on the 2 to +2 s interval surrounding pellet retrieval onset (time 0).

The magnitude of event-aligned LFP activity was quantified using root-mean-square (RMS) amplitude over the 2 to +2 s interval. RMS was calculated for individual trials and then averaged within each animal, yielding one value per hemisphere and learning phase. Learning-related differences were assessed using paired two-sided *t*-tests comparing P1 with P2 and P1 with P3. *P* values were corrected within each hemisphere using the Benjamini–Hochberg false-discovery-rate procedure, and effect sizes were quantified using paired-sample Cohen’s *d_z_*.

#### Power spectral density analysis

Power spectral density (PSD) was estimated separately for the contralateral and ipsilateral RFA using Welch’s method [29]. SPR-aligned LFP activity was analyzed from 2 to +2 s relative to pellet retrieval onset. Spectra were computed for individual trials and channels using 1-s Welch segments over 1–100 Hz and subsequently averaged across channels within each hemisphere. Up to 30 trials per learning phase were included for each animal. For visualization, spectra were averaged first across trials within animals and then across animals, and power was expressed in decibels as 10 log_10_(power).

Learning-related spectral differences were assessed using hierarchical cluster-based permutation tests that accounted for the nesting of trials within animals [30]. Contiguous frequency clusters were tested separately for P1 versus P2 and P1 versus P3 in each hemisphere using 5,000 permutations, with cluster-level *P <* 0.05 considered significant. Analyses were additionally repeated using successful trials only to determine whether learning-related spectral differences were preserved independent of changes in retrieval outcome across learning.

#### Time–frequency analysis

Time–frequency representations (TFRs) of SPR-related LFP activity were computed separately for the contralateral and ipsilateral RFA using complex Morlet wavelets implemented in MNE-Python [28]. Power was estimated from 2 to +1.5 s relative to pellet retrieval onset at 50 geometrically spaced frequencies between 2 and 100 Hz. Wavelet length increased from 3 to 12 cycles across the frequency range. Up to 30 trials per learning phase were included for each animal.

Time–frequency power was calculated for individual trials and channels and normalized to the 3 to 2 s pre-retrieval baseline as 10 log_10_(*P/P*_baseline_). Normalized power was then averaged across channels within each hemisphere. For visualization, trial-level representations were averaged first within animals and subsequently across animals.

TFRs from P1 were compared with those from P2 and P3 separately for each hemisphere using hierarchical cluster-based permutation tests that accounted for the nesting of trials within animals. Adjacent time–frequency samples exceeding the cluster-forming threshold were grouped into clusters, and cluster-level significance was evaluated using 1,000 permutations. Clusters with permutation-based *P <* 0.05 were considered significant. The same comparisons were repeated using successful retrieval trials only.

#### Single-unit firing-rate analysis

Single-unit activity was aligned to pellet retrieval onset and analyzed from 2 to +2 s, with the 3 to 2 s interval used as baseline. Firing-rate time courses were estimated using a 200-ms sliding window advanced in 20-ms steps. Up to 30 trials per learning phase were included for each animal. For each unit and phase, the mean baseline firing rate was subtracted from the event-aligned firing-rate trace to obtain Δ firing rate.

For visualization, firing-rate traces were z-scored within units and displayed separately for P1, P2 and P3 in the contralateral and ipsilateral RFA. Units were ordered independently within each phase according to the timing of their peak firing rate. Z-scored activity was used only for visualization; all statistical analyses were performed on firing rates expressed in hertz.

Within each learning phase, Δ firing rates were tested against zero using two-sided cluster-based permutation tests across units. Adjacent time points exceeding a cluster-forming threshold of *P <* 0.05 were grouped into clusters, and cluster-level significance was evaluated using 5,000 permutations. P1 was additionally compared with P2 and P3 using paired cluster-based permutation tests restricted to units identified across the corresponding phases. Baseline firing rates were compared between phases using two-sided paired *t*-tests, with Benjamini–Hochberg false-discovery-rate correction applied separately within each hemisphere. The same analyses were repeated using successful retrieval trials only.

#### Classification and longitudinal tracking of SPR-responsive units

Units were classified separately at each learning phase according to their firing-rate modulation relative to the pre-retrieval baseline. Using the cluster-based comparison against zero described above, units showing significant increases only were classified as increased, those showing significant decreases only as decreased, those exhibiting both increases and decreases at different time points as mixed, and those showing neither as non-responsive. Decreased units were subsequently referred to as suppression-expressing units.

Response-type proportions were quantified separately for the contralateral and ipsilateral RFA at P1, P2 and P3. For analyses focused on suppression, units were additionally grouped as decreased or non-decreased.

Because recordings from the same electrode channels were jointly sorted across P1, P2 and P3, putative units could be followed across learning phases. Longitudinal response-type transitions were quantified for units identified in both phases of each comparison. Transition matrices were constructed separately for each hemisphere for P1–P2 and P1–P3, with rows representing the response class at P1 and columns representing the corresponding class at the later phase. Matrix entries were expressed as row-normalized percentages.

#### Waveform-based phenotyping of suppression-expressing units

To examine whether SPR-related suppression was preferentially expressed by neurons with distinct extracellular waveform properties, trough-to-peak duration and waveform half-width were quantified for all quality-controlled units. Extracellular spike duration and related waveform features have been widely used to distinguish broad- and narrow-spiking cortical populations [31, 32]. Units with incomplete waveform measurements were excluded.

Broad- and narrow-spiking phenotypes were classified from trough-to-peak duration using Gaussian mixture modeling, with model order selected by the Akaike information criterion, following established waveform-based classification approaches [33]. The component with the shortest mean trough-to-peak duration was designated as narrow-spiking, and the boundary separating narrow-from broad-spiking units was defined by the intersection of its probability density with that of the remaining components. When only a single component was supported, no broad/narrow classification was assigned. Waveform phenotype was defined once for each unit and retained across learning phases.

Waveform phenotype was then related to the phase-specific SPR-response classifications described above. For each hemisphere and learning phase, we quantified the broad- and narrow-spiking composition of suppression-expressing units and the prevalence of suppression within each waveform class. Associations between waveform phenotype and suppression were assessed using Fisher’s exact tests, with Benjamini– Hochberg false-discovery-rate correction applied across learning phases separately within each hemisphere.

#### State space analyses

##### Neural state trajectories

Population activity was analyzed separately for each animal and hemisphere using a state-space approach based on principal component analysis (PCA), adapted from Zheng et al. [34]. Trial-resolved firing-rate activity from P1, P2 and P3 was concatenated across trials and time points, and PCA was applied to define a common state space spanning all learning phases. The minimum number of principal components (PCs) required to explain at least 99% of the cumulative variance was retained. Up to 30 trials per learning phase were included. For visualization, trial trajectories were projected into the common PCA space and averaged within each learning phase.

##### Multidimensional distance

Population neural state displacement was quantified using multidimensional distance (MDD), adapted from Zheng et al. [34], and defined as the Euclidean distance between the population neural state at each time point and the state of the same trial at −2 s in the retained PC space:

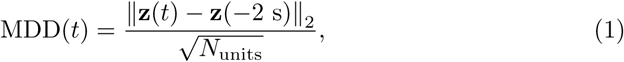

where **z**(*t*) denotes the population neural state in PC space and *N_units_* is the number of units contributing to the population. MDD was normalized by 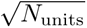 to reduce its dependence on population size.

##### Pre-retrieval MDD

Two trial-level measures were derived from the MDD trajectory. MDD at pellet retrieval onset (*MDD at 0*) quantified the population neural state displacement at time 0, whereas pre-retrieval MDD area under the curve (*Pre-AUC*) quantified the cumulative population neural state displacement over the 2 to 0 s interval. Both measures were calculated for individual trials and examined across learning phases for all trials and separately for successful and failed retrievals.

#### Neural predictors of retrieval success

Trial-level associations between RFA activity and retrieval outcome were assessed using generalized estimating equations (GEE) [35]. Models were fitted separately for the contralateral and ipsilateral RFA with a binomial distribution and logit link, animal identity as the clustering variable and an exchangeable working correlation structure to account for repeated trials within animals.

##### Pre-retrieval firing suppression

For each trial, baseline-corrected firing-rate activity was averaged over the 2 to 0 s interval preceding pellet retrieval. Suppression was defined as the negative of this value, such that larger values indicated stronger pre-retrieval firing suppression. Suppression was standardized within each hemisphere before model fitting. Its association with retrieval success was estimated while accounting for learning phase using

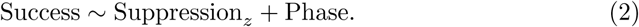

A second model including the suppression-by-phase interaction was used to test whether this association differed across P1, P2 and P3:

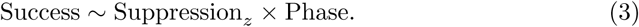

Phase-specific effects and planned P1–P2 and P1–P3 contrasts were obtained from linear contrasts of the fitted model. Regression coefficients were exponentiated to obtain odds ratios (ORs) and 95% confidence intervals, with ORs corresponding to a 1-s.d. increase in suppression. The overall interaction was assessed using a two-degree-of-freedom Wald test, and Benjamini–Hochberg false-discovery-rate correction was applied separately to the three phase-specific tests and the two planned contrasts.

##### Population neural state engagement

The same GEE framework was applied separately to MDD at pellet retrieval onset (*MDD at 0*) and pre-retrieval MDD area under the curve (*Pre-AUC*). Each metric was standardized within hemisphere before analysis. Overall associations with retrieval success were estimated using

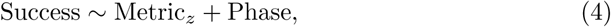

where Metric*_z_* represented either standardized MDD at 0 or standardized Pre-AUC. Phase dependence was tested using

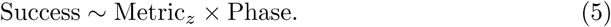

For each metric and hemisphere, phase-specific ORs and planned P1–P2 and P1–P3 contrasts were obtained from the fitted interaction model. Omnibus interactions were evaluated using two-degree-of-freedom Wald tests, and Benjamini–Hochberg false-discovery-rate correction was applied separately to the phase-specific tests and planned contrasts.

### Statistical analysis

Statistical analyses were performed in Python 3.10.20 using SciPy 1.15.3 [36] and Statsmodels 0.14.6 [37]. All tests were two-sided unless otherwise specified, with statistical significance defined as *P <* 0.05. Data are reported as mean ± s.e.m. unless stated otherwise.

Individual animals were treated as independent experimental units for animal-level analyses, with paired tests used for longitudinal comparisons across learning phases. Trial-level analyses accounted for repeated observations within animals using hierarchical permutation tests or generalized estimating equations, as described in the corresponding sections. Longitudinal single-unit comparisons were restricted to putative units identified across the relevant learning phases.

Multiple comparisons were controlled using the Benjamini–Hochberg false-discovery-rate procedure where indicated [38]. Cluster-based permutation tests were used for time-, frequency- and time–frequency-resolved analyses [30, 39]. Effect sizes for paired animal-level comparisons were reported as Cohen’s *d_z_*, and associations with retrieval success were summarized as odds ratios with 95% confidence intervals. Exact sample sizes, statistical tests and adjusted or unadjusted *P* values are reported in the corresponding Results and figure legends.

## Supplementary information

Fig. S1 and its legend in a PDF

Fig. S2 and its legend in a PDF

## Acknowledgements

This work was supported by the Brain Science and Brain-like Intelligence Technology-National Science and Technology Major Project (2022ZD0208900) and the National Natural Science Foundation of China (U22A20293, 62306120).

## Declarations

The authors declare no competing interests.

## Author contributions

Conceptualization, Yuanqing Li and Ying Guo; methodology, Di Chen; Formal analysis, Di Chen and Fang Jia; investigation, Fang Jia; writing – original draft, Di Chen; writing – review & editing, Di Chen and Qiyun Huang; funding acquisition, Yuanqing Li; resources, Ying Guo; supervision, Yuanqing Li and Ying Guo.

## Resource availability

### Lead contact

Requests for further information and resources should be directed to and will be fulfilled by the lead contact, Di Chen.

### Materials availability

This study did not generate new, unique reagents.

### Data and code availability

The data supporting the findings of this study are available from the lead contact upon reasonable request. All custom code used for data processing, analysis, and figure generation will be made publicly available in an online repository upon publication.

## Notes

### Competing Interest Statement

The authors have declared no competing interest.

